# Redox signaling by hydrogen peroxide modulates axonal microtubule organization and induces a specific phosphorylation signature of microtubule proteins distinct from distress

**DOI:** 10.1101/2024.07.01.601594

**Authors:** Christian Conze, Nataliya I. Trushina, Nanci Monteiro-Abreu, Daniel Villar Romero, Eike Wienbeuker, Anna-Sophie Schwarze, Michael Holtmannspötter, Lidia Bakota, Roland Brandt

**Affiliations:** Department of Neurobiology, School of Biology/Chemistry, Osnabrück University, Germany; Center for Cellular Nanoanalytics, Osnabrück University, Germany; Institute of Cognitive Science, Osnabrück University, Germany

**Keywords:** microtubules, tau, axon, redox signalling, hydrogen peroxide

## Abstract

Many life processes are regulated by physiological redox signals, referred to as oxidative eustress. However, excessive oxidative stress can damage biomolecules and contribute to disease. The neuronal microtubule system is critically involved in axon homeostasis, regulation of axonal transport, and neurodegenerative processes. However, whether and how physiological redox signals affect axonal microtubules is largely unknown. Using live cell imaging and super-resolution microscopy, we show that subtoxic concentrations of the central redox metabolite hydrogen peroxide increase axonal microtubule dynamics, alter the structure of the axonal microtubule array, and affect the efficiency of axonal transport. We report that the mitochondria-targeting antioxidant SkQ1 and the microtubule stabilizer EpoD abolish the increase in microtubule dynamics. We found that oxidative eustress and distress specifically modulate the phosphorylation state of the microtubule system and induce a largely non-overlapping phosphorylation pattern of MAP1B as the main target. Cell-wide phosphoproteome analysis revealed that different signaling pathways are inversely activated by oxidative eustress and distress. Signaling via casein kinase (CK2) and pyruvate dehydrogenase kinases (PDK) is activated during eustress and signaling via mammalian target of rapamycin (mTOR) and serum/glucocorticoid-regulated protein kinase (SGK) is activated during distress. The results suggest that the redox metabolite and second messenger hydrogen peroxide induces rapid and local reorganization of the microtubule array in response to mitochondrial activity or as a messenger from neighboring cells by activating specific signaling cascades.

## INTRODUCTION

Microtubules are a crucial filament system involved in virtually all aspects of a neuron’s life. In the axon, they have a characteristic organization consisting of an array of relatively short microtubules, the plus-end of which is oriented towards the distal tip (Penazzi et al., 2016a). Such organization is thought to be critical for efficient axonal transport, in which cargo is carried to the distal axon over long distances. Although axonal microtubules are relatively stable, they still show considerable dynamics, with the plus ends of the microtubules exhibiting stochastic changes between growth and shrinkage, referred to as dynamic instability. Microtubule stability and dynamics are regulated by a variety of tubulin- and microtubule-interacting proteins, including microtubule nucleators, microtubule-binding proteins, end-binding proteins, tubulin-sequestering proteins and microtubule-severing proteins. Further complexity arises from the presence of a variety of different tubulin isoforms, where humans have eight α− and nine β−tubulin genes, most of which are expressed in neurons (Uhlen et al., 2015). Many of the components of the microtubule system have extensive posttranslational modifications (PTMs) that modulate their activity. The best studied PTM of the microtubule system is phosphorylation, and increased phosphorylation of specific serine and threonine residues is known to influence the interaction of neuronal microtubule-associated proteins (MAPs) such as MAP1B and tau, with microtubules. Of disease-related importance is the modulation of the interaction of the MAP tau with axonal microtubules, as tau is hyperphosphorylated in Alzheimer’s disease and other tauopathies, which largely reduces its interaction with microtubules (Arendt et al., 2016).

Oxidative stress is associated with many neurodegenerative diseases, and increased levels of reactive oxygen species (ROS) may play a role in triggering axonal degeneration and microtubule disassembly (Pratico et al., 2002). High levels of ROS form additional free radical compounds that have the potential to damage lipids, proteins, and nucleic acids over time (Vincent et al., 2009). Indeed, high levels of ROS have been shown to inhibit axonal transport (Fang et al., 2012) and induce axonal degeneration, as evidenced by the morphological features of axon beading and fragmentation (Fukui et al., 2011). However, more recently it has become clear that moderately elevated ROS levels can act as a crucial physiological mediator of many cellular processes (Sinenko et al., 2021). This led to the distinction of “oxidative distress” as a mechanism leading to pathology and “oxidative eustress” as an important physiological regulator of intrinsic signalling pathways (Wilson et al., 2018).

The most common ROS members in mediating oxidative eustress are the superoxide anion, the hydroxyl radical, and hydrogen peroxide. Because these ROS species act as relatively short-lived second messengers, they are able to regulate intrinsic signalling pathways (Wilson et al., 2018). Hydrogen peroxide was found to be the most important redox metabolite involved in signal transduction and redox regulation. It is an electrically neutral molecule and is chemically more stable than the other members of the ROS family. As a messenger molecule, hydrogen peroxide diffuses through cells and tissues to trigger immediate cellular effects that link redox biology with regulation through phosphorylation and dephosphorylation (Sies, 2017). Physiological hydrogen peroxide levels promote the establishment of neuronal polarity, neurite growth and axon specification (Wilson et al., 2015). While all of these processes critically depend on the regulation of microtubule polymerization, the mechanisms of how hydrogen peroxide influences axonal microtubules and microtubule-dependent functions under oxidative eustress conditions are largely unknown.

Here, we determined the effect of subtoxic concentrations of hydrogen peroxide, reflecting oxidative eustress, on axonal microtubule dynamics using quantitative live-cell imaging. We analyzed changes in the structure of the axonal microtubule array using single-molecule localization microscopy and algorithm-based reconstruction of microtubules. We examined possible functional microtubule-dependent consequences by tracking individual APP vesicles and determining the tau-microtubule interaction using fluorescence-decay after photoactivation (FDAP) experiments. To identify downstream targets of increased ROS, we performed proteomic and phosphoproteomic analysis of differentiated neurons treated with subtoxic hydrogen peroxide and compared them with arsenite treatment as an inducer of oxidative distress-mediated toxicity.

## MATERIALS AND METHODS

### Materials

Unless otherwise stated, chemicals and cell culture material were purchased from Sarstedt (Nümbrecht, Germany), Sigma-Aldrich (Deisenhofen, Germany), and Thermo-Fisher Scientific (Waltham, USA). The microtubule-targeting agent epothilone D (EpoD) was a kind gift from Amos Smith 3rd (University of Pennsylvania) and was prepared as previously described (Lee et al., 2001; Rivkin et al., 2004). The purity of the compound was > 95%, as determined by LC-MS and NMR analyses. The spectroscopic properties were identical to those reported in the literature. The mitochondria-targeted antioxidant SkQ1, and TPP, which lacks the antioxidant quinone moiety, were provided by Maxim Skulachev (Mitotech S.A., Luxembourg).

### Cell culture and transfections

PC12 cells were cultured in serum-DMEM and transfections were performed with Lipofectamine 2000 (Thermo-Fisher Scientific, USA) as previously described (Fath et al., 2002). Expression plasmids for PAGFP-α-Tubulin and mEGFP-α-Tubulin have been described previously (Conze et al., 2022b). The expression plasmid PAGFP-htau441wt (Gauthier-Kemper et al., 2011) was used to express the longest CNS-isoform of tau. pEGFP-n1-APP was obtained from Zita Balklava and Thomas Wassmer (Addgene plasmid #69924; http://n2t.net/addgene:69924; RRID: Addgene_69924).

### Metabolic activity and cytotoxicity profiling using a combined LDH and MTT Assay

PC12 cells were cultured in 96-well plates at 1×10^4^ cells/well in serum-reduced medium supplemented with 100 ng/ml 7S mouse NGF. Cells were incubated for 48 hours to initiate neuronal differentiation. Concentration-response profiling of hydrogen peroxide (H2O2) was performed with four triplicates and six concentrations ranging from 50-900 µM. The respective H2O2 concentration was added to each well 3 hours before the measurement. Designated wells for controls were untreated for negative control or a total cell lysis was induced by the addition of 1% Triton-X for a positive control. For the LDH assay, 50 µl medium from each well of the assay plate was transferred to a separate 96-well plate. For quantification of the LDH release, 50 µl of LDH reagent (4 mM iodonitrotetrazolium chloride (INT), 6.4 mM beta-nicotinamide adenine dinucleotide sodium salt (NAD), 320 mM lithium lactate, 150 mM of 1- methoxyphenazine methosulfate (MPMS) in 0.2 M Tris-HCl buffer, pH 8.2) was added. The plate was shaken for 10 seconds and incubated in the dark for several minutes. Absorbance was measured at 490 nm using a Thermomax Microplate Reader operated with SoftMaxPro Version

1.1 (Molecular Devices Corp., Sunnyvale, CA, U.S.A.). LDH release measurements were normalized to the optical densities of the positive control wells. For the MTT assay, MTT reagent (3,(4,5.dimethylthiazol-2-yl)2,5-diphenyltetrazolium bromide) at a final concentration of 1 mg/ml MTT was added to the wells in the remaining assay plate. The cells were incubated for a further 2 hours in the cell culture incubator before the reaction was stopped by adding 50 μl of lysis buffer (20% (wt/vol) sodium dodecyl sulfate in 1:1 (vol/vol) N,N- dimethylformamide/water, pH 4.7). After overnight incubation at 37 °C, the optical densities of the formazan product were determined at 570 nm. MTT conversion measurements were normalized to the optical densities of the negative control wells.

### Detection of intracellular ROS levels

Intracellular ROS levels were assessed using the Cellular ROS Assay Kit Orange (abcam, ab186028, Cambridge, UK). PC12 cells (2×10^4^ cells/well) were seeded in 60 μl of serum-reduced DMEM supplemented with 100 ng/ml 7S mouse NGF in a 96-well plate. To measure the effect of SkQ1, 30µl of NGF containing serum-reduced medium supplemented with 0.2 µM SkQ1 or carrier control (0.2% EtOH) were added after 24 hours. After a further 24 hours of incubation, the cells were treated with the respective H2O2 concentration (150 µM and 450 µM) for 3 hours. A total of six independent experiments were conducted. Intracellular ROS levels were determined according to the manufacturer’s instructions. Briefly, cells were incubated with 100 µl of ROS Orange Working solution for 60 min. Changes in fluorescence intensity were measured with the FLUOstar Optima (BMG Labtechnologies, Ortenberg, Germany) at Ex/Em = 540/570 nm. The fluorescence intensities were normalized to the respective negative control values after background subtraction.

### Live-cell imaging

Fluorescence decay after photoactivation (FDAP) experiments with PAGFP-α-tubulin- or PAGFP-tau-expressing cells were performed essentially as described previously (Niewidok et al., 2016). Briefly, PC12 cells were plated on 35-mm poly-L-lysine and collagen-coated glass-bottom culture dishes (MatTek, USA). After transfection, PC12 cells were neuronally differentiated by media exchange with DMEM with 1% (vol/vol) serum containing 100 ng/ml 7S mouse NGF (Alomone Laboratories, Germany). Cultivation was continued for 4 days with medium exchange to DMEM with 1% (vol/vol) serum containing NGF and without phenol red one day prior to live imaging. Live imaging of PC12 cells for photoactivation experiments was performed using a Nikon Eclipse Ti2-E laser scanning microscope (Nikon, Japan) equipped with a LU-N4 laser unit with 488-nm and 405-nm lasers and a Fluor 60× ultraviolet-corrected objective lens (NA 1.4) enclosed in an incubation chamber at 37°C and 5% CO2. Automated image acquisition of PAGFP-tubulin- or PAGFP-tau-expressing cells after photoactivation was performed essentially as described previously (Igaev et al., 2014). Briefly, photoactivation of a 6 μm long neurite segment was performed with a 405-nm laser. A set of consecutive image series (time stacks) was created at a frequency of 1 frame/s and 112 frames were collected per activated cell with a resolution of 256×256 pixels.

Live-cell imaging to analyze APP transport in PC12 cells was performed essentially as previously described (Conze et al., 2022a) using a Zeiss Cell Observer Z1 (Zeiss, Germany) with high-speed confocal imaging using the CSU-X1 spinning disc technology from Yokogawa, equipped with an optically pumped 488-nm semiconductor laser surrounded by a sample incubation chamber controlled by a Zeiss TempModule S1. pH stability was maintained by adding 30 mM HEPES buffer to the cell culture medium before image acquisition. For image capture of eGFP-APP vesicle tracking, the 488-nm laser and Alpha Plan-Apochromat ×63 (NA 1.46) objective were used. An image series (time stack) of 60 s was acquired at a speed of five images per second, captured with a Hamamatsu ORCA flash V3 and 2×2 binning at a resolution of 640×640 pixels (pixel size 172 nm). The image series were deconvolved with the software Huygens Remote Manager v3.5 (Scientific Volume Imaging B.V., Hilversum, Netherlands) using a theoretical PSF and a classical maximum likelihood estimation as the deconvolution algorithm with 20 iterations and a quality criterion of 0.05.

Treatment of PC12 cells with 150 µM H2O2 was performed 3 hours before live imaging. To assess the effect of the antioxidant SkQ1 in FDAP experiments, cells were treated with 0.2 µM of the compound 24 hours before H2O2 addition. To determine the effect of H2O2 treatment on stabilized MTs by epothilone D (EpoD) in FDAP experiments as well as APP vesicle tracking, cells were treated with 1 nM EpoD or its carrier control (0.01% DMSO) one hour before H2O2 treatment.

### FDAP data analysis

A reaction-diffusion model was used to determine the association rate k^*^on and the dissociation rate koff constant of tubulin or tau binding, as described previously (Igaev et al., 2014). Processing and analysis of individual FDAP curves was performed as previously described (Niewidok et al., 2016) using a custom C-based tool called cFDAP. The fitting procedure was used to obtain k^*^on and koff from each individual FDAP curve, while the χ2 value was used as an indicator of the goodness of fit of the model function. Effective diffusion constants were obtained by fitting fluorescence decay data from photoactivation experiments using a one-dimensional diffusion model function for FDAP, as previously described (Weissmann et al., 2009).

### Tracking and quantifying axonal transport

APP transport was tracked and analyzed using Imaris v 9.2 (Bitplane, Oxford Instruments). Vesicles were detected by the Gaussian-filtered intensity of their signal within a diameter of 500 nm around the respective signal peak. Vesicle transport was tracked by an autoregressive motion algorithm with a maximum distance of a future position of the signal of 1 µm and a maximum gap size of 2 frames (400 ms). To determine the direction of the tracked vesicles, a reference point at the transition from the cell body to the neurite was used, while only the tracks that spanned a period of at least 3 s (15 frames) were considered for further analysis. Quantified transport parameters were exported to spreadsheets (Excel, Microsoft Corporation, USA) and further processed using TIBCO Spotfire Data Analysis Software (TIBCO Software Inc., USA). Vesicles that exceeded a displacement of more than 0.75 µm over the observation time of 60 s were designated as mobile, otherwise as stationary.

### Super-resolution microscopy with DNA-PAINT

For super-resolution microscopy, PC12 cells were transfected with mEGFP-α-Tubulin and neuronally differentiated for 4 days as described above. Fixation, immunostaining, and super-resolution microscopy with DNA-PAINT were performed essentially as previously described (Conze et al., 2022b). Briefly, PC12 cells were processed using a combined NP-40 permeabilization-fixation protocol, which removes membranes and cytosolic components but preserves cytoskeletal structures and associated proteins (Brandt et al., 1995). Immunostaining used an anti-GFP nanobody obtained from the Massive-Tag-Q Anti-GFP DNA-PAINT kit (Massive Photonics GmbH, Gräfelfing, Germany). Just before imaging, cells were washed three times with PBS and incubated with a 1:1000 dilution of 50 nm gold nanorods (Nanopartz, E12- 50-600-25) that function as fiducial markers during image acquisition. Afterwards, cells were washed three times with PBS before an imaging buffer (Massive Photonics) supplemented with 250 pM Cy3b-conjugated imager DNA-strand, complementary to the DNA strand attached to the anti-GFP nanobody, was added. Cells were imaged by total internal reflection fluorescence (TIRF) microscopy using an inverted microscope frame (Olympus IX-81) equipped with a motorized quad-line TIR illumination condenser (cellTIRF-4-Line, Olympus) and a motorized xy-stage (Märzhäuser Scan IM 120x80). Three-dimensional single-molecule localization was achieved by astigmatic imaging using a cylindrical lens (Olympus) implemented directly in front of the filter wheel.

### Post-processing of DNA-PAINT datasets

Raw data sets were processed using the "Picasso" software package (Schnitzbauer et al., 2017), https://github.com/jungmannlab/picasso). At the beginning of each imaging session, a z-stack of immobilized fluorescent TetraSpeck™ microspheres with a diameter of 100 nm (Invitrogen, T7279) was acquired. This was done in the respective imaging buffer with a step size of 10 nm using a piezo z-stage (NanoScanZ, NZ100, Prior Scientific). This is a necessity for the calibration of astigmatic PSFs, which is a prerequisite for three-dimensional single-molecule localization microscopy. This z-stack was analyzed using “Picasso: Localize” with parameters, that allow the identification of single beads in each frame. The photon conversion parameters were set as follows: EM Gain: 1, Baseline: 400, Sensitivity 0.46, Quantum Efficiency: 0.80 and pixel size: 130 nm. A calibration file was generated with the "Calibrate 3D" function of "Picasso localize". This file, as well as the same photon conversion parameters, were used for image processing of raw sample files. The Min. Net. Gradient was adjusted to remove non-specifically bound imager strands or other background signals to filter for localizations with the highest signal intensities. Single-molecule localizations were fitted with a Gaussian least-square fit. For 3D localization, the magnification factor was set to 1.0. Processed data sets were opened with "Picasso: Render" and a drift correction with cross-correlation was performed, followed by a correction using the fiducials. The localizations of these datasets were then exported for the ImageJ plugin ThunderSTORM for 3D rendering to obtain image-stacks with a well-defined voxel size set to 26 nm×26 nm×25 nm based on the overall axial resolution of 25 nm (Ovesny et al., 2014).

### SIFNE analysis of post-processed DNA-PAINT data

Computational analysis and quantification of the microtubule array in axon-like processes of PC12 cells was performed using the open-source SIFNE (SMLM image filament extractor) software package (Zhang et al., 2017). The MATLAB-based tool involves the iterative extraction of the filamentous structures from the image data set and the subsequent identification and assignment of the detected filaments. Since the axial distance between microtubules in PC12 neurites is about 70 nm (Jacobs and Stevens, 1986), we chose optical sections with a thickness of 150 nm for filament extraction by SIFNE. This allowed us to use sections where the resolution was highest while avoiding overlapping microtubules from another plane that would confound our statistics. The region of interest (ROI) in axon-like processes of PC12 cells was set to the shaft of the neurite. For image enhancement using line and orientation filter transformation algorithms (LFT and OFT), a radius of 10 pixels with 40 rotations of the scan line segment was used. Segmentation was performed using SIFNE’s automatic thresholding function based on Otsu’s method. For creating a pool of minimal linear filament fragments, areas of filament connections were removed by a local area around each connection of 2×2 pixels. To recover unrecognized linear structures, iterative extraction of the filaments was performed twice, followed by registration of the propagation direction of each filament tip. The grouping and analysis of the detected filaments was carried out with a pixel size of 26 nm and a maximum curvature of 1 rad/µm. The search angle and radius were set to 60° and 40 pixels, respectively. The allowable orientation difference between the endpoints was set to 60°. The maximum allowable angle difference and endpoint gap vector were set to 60° and 30°, respectively. The weights for similarity and continuity conditions during the scoring calculations were set to 1. Due to the high complexity of the cytoskeletal network, fragment overlap was not allowed as suggested by (Zhang et al., 2017). For sorting composite filaments, the minimum filament length was set to 15 pixels corresponding to 390 nm, while ungrouped filaments were left in the data set.

### Proteome and phosphoproteome analysis

PC12 cells were neuronally differentiated by culture in serum-reduced DMEM with 100 ng/ml 7 S mouse NGF for 4 days and treated with 150 µM H2O2 for 3 hours, 0.5 mM arsenite for 20 min, or left as respective controls. Proteome and phosphoproteome analysis were essentially performed as described previously (Pinzi et al., 2024). Briefly, cells were incubated in lysis buffer (8M urea in 50mM Tris/HCl, pH 7.8) supplemented with Phos-Stop tablets (Roche Diagnostics GmbH, Germany), sonicated and cleared by centrifugation. Protein concentrations were determined using Pierce™ BCA Protein Assay (Thermo Fisher Scientific, USA). 0.2 μg/μl α-casein was added to a protein amount of 1.2 mg, and reduction and alkylation were carried out in lysis buffer containing 15 mM iodoacetamide and 5 mM DL-dithiothreitol. The proteins were digested with trypsin/Lys-C Mix (Promega Corporation, USA), and 10 μg of each sample was used for proteomic analysis. For phosphoenrichment, the remaining samples were desalted using Sep-Pak® Classic C18 cartridges (Waters, Ireland). The eluate was lyophilized and enriched with a High-Select™ TiO2 Phosphopeptide Enrichment Kit (Thermo Fisher Scientific, USA). For proteome and phosphoproteome analysis, samples were collected from a PepMap C18 easy spray column (Thermo Fisher Scientific, USA). MS analysis was performed as previously described (Schoppe et al., 2020). The *.raw data files were analyzed using PEAKS Online software (Bioinformatic Solutions Inc, Canada). PEAKS Q (de-novo-assisted quantification) analysis was used for data refinement with mass correction, de novo sequencing and de novo-assisted database search, and subsequent label-free quantification. The search engine was applied to Rattus norvegicus *.fasta databases. MS/MS searches were performed using a mass tolerance of 10 ppm parent ions and a mass tolerance of 0.2 Da fragments. Trypsin with up to two missing cleavage sites was selected as the cleavage enzyme. The carbamidomethylation modification was chosen as the fixed modification and the oxidation of methionine, the acetylation of lysine and the phosphorylation of serine, threonine and tyrosine as variable modifications. A maximum of three variable modifications were allowed per peptide. Normalization to the total ion current level was performed for each sample. The ANOVA test was used to calculate the significance level for each protein, outliers were removed, and the top three peptides were used to quantify the protein signal where possible. The peptide identification was considered valid at a false detection rate of 1% (q-value < 0.001) (maximum delta Cn of the percolator was 0.05). The minimum length of acceptable identified peptides was set to six amino acids. Each condition was analyzed in triplicate. All proteins were assigned their gene symbol via the Uniprot knowledge database (http://www.uniprot.org/).

### Bioinformatic analysis

The components of the microtubule system highlighted on the volcano plots were based on the classification of the different functional groups of the microtubule system in (Trushina et al., 2019b): structure proteins (α- and β-tubulins), nucleators (γ-tubulins and γ-tubulin complex proteins (GCPs)), MT-binding proteins (MAP1A, MAP1B, MAP1S, MAP2, tau (encoded by the MAPT gene), MAP4, MAP6 (STOP) and MAP7 (ensconsin)), tubulin-sequestering proteins (stathmins, encoded by STMN1, STMN2 (SCG10), STMN3 (SCLIP), STMN4 (RB3)), end-binding proteins (EB1, EB2, EB3 (encoded by MAPRE1, MAPRE2, MAPRE3, respectively), CLASP1 and CLASP2), MT-severing proteins (P60-katanin (encoded by KATNA1 and KATNB1), fidgetin (FIGN) and spastin (SPAST)). Gene Ontology (GO) term enrichment analysis was performed to identify overrepresented Molecular Function (MF) and Cellular Component (CC) categories using EnrichR (Xie et al., 2021). All differentially expressed proteins and all proteins with upregulated phosphosites were used to create gene sets for enrichment analysis. The 10 highest-scoring GO-terms for molecular function and cellular component were displayed. Gene Ontology (GO) regarding molecular function and cellular component was used to construct of protein classifications. Kinase enrichment analysis (KEA) was performed using the curated kinase-substrate database KEA2 (Lachmann and Ma’ayan, 2009). The gene symbols with phosphorylated sites of differentially phosphorylated proteins were used to infer upstream kinases whose putative substrates are overrepresented. To identify the most important functional pathways upon treatment with hydrogen peroxide or arsenite, significantly enriched kinases were compared.

### Statistical analysis

Statistical analysis was performed using GraphPad Prism v8.0.1 (GraphPad Software, USA). All datasets were tested for normality using the D’Agostino-Pearson and Shapiro-Wilk tests. When necessary, datasets were log transformed to allow further statistical testing. Statistical outliers were identified using the ROUT method. Homogeneity was assessed using the Levene test. An unpaired two-tailed t-test was used to compare two datasets. In cases of unequal variances, Welch’s correction was applied. To compare more than two data sets, one-way ANOVA was performed followed by Dunnet post-hoc test. All statistical values are expressed as mean ± SEM.

## RESULTS

### Subtoxic concentrations of hydrogen peroxide decrease microtubule polymer in axon-like processes by decreasing kon and increasing koff rates of microtubule polymerization

The redox metabolite hydrogen peroxide is known to act as a messenger molecule and diffuses through cells and tissues (Sies, 2017). While high concentrations of hydrogen peroxide are considered toxic (“oxidative distress”), low levels of hydrogen peroxide have been shown to promote neuronal development and axon specification (Wilson et al., 2015), thus positively modulating cellular functions through a process called “oxidative eustress” (Wilson et al., 2018). Therefore, we first determined the level of subtoxic concentrations of hydrogen peroxide in model neurons, which are likely to induce eustress mechanisms in the cells.

We used a combined MTT and LDH assay to determine the metabolic activity and cytotoxicity profile of exogenously added hydrogen peroxide on differentiated neuronal cells. We observed that a short (3 hour) treatment with 150 µM hydrogen peroxide affected neither MTT conversion, as a measure of metabolic activity, nor LDH release, as a measure of toxicity (Fig. 1A). In contrast, an increase to 225 µM hydrogen peroxide induced both a decrease in MTT conversion and an increase in LDH release, both of which became highly significant at 450 µM. Treatment with subtoxic 150 µM hydrogen peroxide increased intracellular ROS level by ∼30%, while toxic 450 µM hydrogen peroxide resulted in an increase of ∼70% (Fig. 1B).

**Fig. 1.**
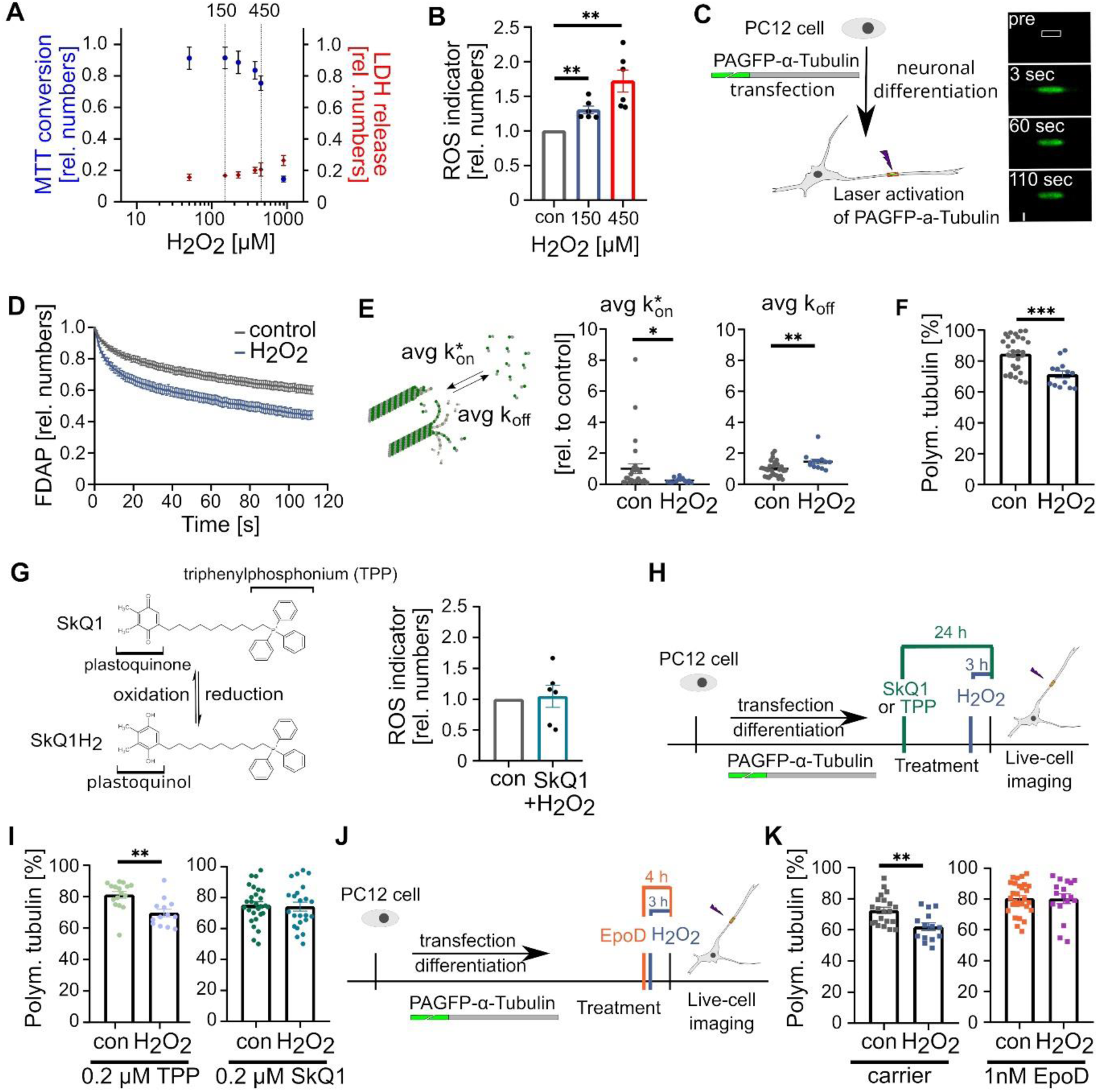
Subtoxic concentrations of hydrogen peroxide decrease microtubule polymer in axon-like processes by decreasing kon and increasing koff rates of microtubule polymerization. **A.** Combined MTT (blue) and LDH (red) assay showing the effect of 3 h H2O2 exposure on neuronally differentiated PC12 cells. Mean ± SEM of 4 experiments, each carried out in triplicates, are shown. Statistically significant differences as determined by one-way ANOVA followed by a Dunnett post hoc test showed significant differences from a control at concentrations ≥ 225 µM. **B.** Bar graphs showing the increase in intracellular ROS level in response to H2O2. Each data point represents an independent experiment normalized to a control. Mean ± SEM is shown. Statistically significant differences between treated and control cells as determined by one-sample t-tests are indicated (**p<0.01). **C.** Representative time-lapse images of a fluorescence decay after photoactivation (FDAP) experiment in an axon-like process. A 6 µm long segment (white box) in the middle of a process was photoactivated and the fluorescence decay in this area was monitored over time. A schematic representation of the FDAP approach and the expressed construct is shown on the left. **D.** FDAP diagrams after photoactivation of PAGFP-α-Tubulin expressing cells show an increased fluorescence decay after treatment with H2O2. Mean values ± SEM of 29 (control) and 13 (150 µM H2O2) cells are shown. **E.** Scatterplots of the association (k^*^) and dissociation rate constants (k), determined on by modelling the FDAP plots from (D), show that H2O2 decreases k^*^ off and increases koff. Statistically significant differences between treated and control cells, determined by unpaired two-tailed Student’s t-tests, are indicated. *p<0.05, **p<0.01. **F.** The scatterplot of the amount of polymerized tubulin determined from the association and dissociation constants in (E) shows that H2O2 reduces the amount of polymerized tubulin in axon-like processes. Statistically significant differences determined by unpaired two-tailed Student’s t-tests are indicated. ***p<0.001. **G.** A schematic representation of the conversion between the oxidized and reduced forms of SkQ1 as a mitochondria-targeted antioxidant is shown. TPP, which lacks the antioxidant quinone moiety, is indicated. A bar graph is displayed on the right showing that pretreatment with SkQ1 abolished the increase in intracellular ROS levels in response to H2O2. **H.** Schematic representation of the timeline of the experiment to determine the effect of pretreatment with SkQ1 or a control (TPP) on microtubule polymerization. **I.** Scatterplot of the amount of polymerized tubulin with the control (TPP) and SkQ1, showing that pretreatment with SkQ1 prevents H2O2-induced microtubule depolymerization in axon-like processes as determined by FDAP experiments. Shown are mean values ± SEM of 16, 13 (TPP) and 28, 23 (SkQ1) cells for control and H2O2-treatment, respectively. Statistically significant differences determined by unpaired two-tailed Student’s t-tests are indicated. **p<0.01. **J.** Shown is the time course of the experiment to determine the effect of pretreatment with the microtubule-stabilizer EpoD on H2O2-induced microtubule depolymerization. **K.** Scatterplot of the amount of polymerized tubulin showing that pretreatment with EpoD prevents H2O2-induced microtubule depolymerization. Shown are mean values ± SEM of 21, 14 (carrier, 0.01% DMSO) and 29, 17 (1 nM EpoD) cells for control and H2O2-treatment, respectively. Statistically significant differences determined by unpaired two-tailed Student’s t-tests are indicated. **p<0.01.

In further experiments, we therefore decided to focus on treating neuronal cells with the subtoxic concentration of 150 µM hydrogen peroxide, which results in only a moderate increase in intracellular ROS levels, likely reflecting “oxidative eustress”. Because neuronal development and function critically depend on the regulation of microtubule polymerization, we determined the effect of hydrogen peroxide on axonal microtubule dynamics using a previously established live cell imaging approach. The method is based on fluorescence decay after photoactivation (FDAP) measurements of model neurons transfected to express PAGFP-tagged α-tubulin (PAGFP-α-tubulin). Changes in microtubule dynamics were then monitored by FDAP measurements on cells in which the fluorescence of PAGFP was focally activated in the middle of an axon-like process (Fig. 1C). FDAP curves showed that hydrogen peroxide treatment resulted in increased fluorescence decay, indicating a lower proportion of polymerized tubulin (Fig. 1D). Notably, application of a reaction-diffusion model resulted in greatly reduced k^*^, indicating that hydrogen peroxide induces microtubule depolymerization (Fig. 1E). The significant increase in koff indicates higher microtubule dynamics as it reflects a shortened residence time of the tubulin dimers in the polymer. The amount of polymerized tubulin in the axon-like processes was reduced by ∼10% from more than 80% to ∼70% (Fig. 1F).

Taken together, the data indicate that subtoxic concentrations of hydrogen peroxide modulate the properties of the axonal microtubule array by reducing the proportion of microtubule polymer and increasing their dynamics.

### The mitochondria-targeted antioxidant SkQ1 and the microtubule stabilizer epothilone D prevent hydrogen peroxide-induced microtubule depolymerization

A major source of ROS under physiological conditions is mitochondria. Through their NAD^+^ pool, they play a predominant role in defining cellular responses to stress. While the mitochondrial NAD^+^/NADH redox ratio is maintained separately, it appears to be strongly connected to the cytosolic NAD^+^ pool and responsive to changes in its perturbations (Hu et al., 2021). Therefore, we asked whether targeting the redox status of mitochondria would influence hydrogen peroxide-mediated modulation of microtubule dynamics.

To modulate the mitochondrial redox state, we used the mitochondria-targeted antioxidant SkQ1. SkQ1 is a derivative of plastoquinone, a potent antioxidant (Antonenko et al., 2008) and has already been used in preclinical studies to treat cardiovascular and renal diseases (Fedorov et al., 2022). Indeed, treatment with SkQ1 completely abolished the hydrogen-peroxide induced increase in intracellular ROS levels (Fig. 1G). The same was true for the hydrogen peroxide-induced modulation in microtubule polymer, where pretreatment with SkQ1 completely abolished the effect of hydrogen peroxide on microtubule polymer reduction in axon-like processes (Fig. 1H, I). In contrast, TPP lacking the antioxidant quinone moiety had no effect (Fig. 1I) confirming the antioxidant activity of SkQ1 in preserving the state of axonal microtubules.

If the cytosolic redox state indeed modulates axonal microtubule dynamics, thereby leading to reduced microtubule polymer, a microtubule stabilizer may prevent this effect. To test this hypothesis, we used the well-characterized microtubule-targeting agent (MTA) epothilone D (EpoD). This small molecule microtubule stabilizer binds to the β-tubulin subunit on the luminal surface of microtubules, induces tubulin polymerization similar to paclitaxel (Buey et al., 2004), and reduces microtubule-dependent spine loss in an Alzheimer’s disease model already at subnanomolar concentrations (Penazzi et al., 2016b). In fact, treatment with nanomolar EpoD abolished the decrease in microtubule polymer caused by hydrogen peroxide (Fig. 1J, K).

Thus, the data indicate that the mitochondrial redox state modulates microtubule dynamics in the axonal compartment. Furthermore, they show that microtubule-stabilizing drugs override the modulation of microtubule dynamics by the cytosolic redox state.

### Subtoxic hydrogen peroxide modulates the structure of the axonal microtubule array by reducing microtubule mass and increasing microtubule length

Axonal microtubules are not continuous but are present in an array of relatively short, uniformly oriented fragments. In developing mammalian axons, average microtubules are only a few micrometers long (Bray and Bunge, 1981; Yu and Baas, 1994). The question therefore arises as to whether hydrogen peroxide also changes the microtubule arrangement in axons, for example their length distribution or their physical properties.

To determine microtubule organization in axon-like processes, we used single molecule localization microscopy (SMLM) using DNA-PAINT (Point Accumulation in Nanoscale Topography) (Jungmann et al., 2010), followed by algorithm-based filament extraction (Zhang et al., 2017) (Fig. 2A). We have previously validated the approach in axon-like processes of PC12 cells and dendrites of primary neurons (Conze et al., 2022b). As expected from the FDAP data, hydrogen peroxide resulted in reduced microtubule polymer, as evidenced by lower microtubule mass and density (Fig. 2B, left). Interestingly, hydrogen peroxide led to an increase in mean microtubule length (Fig. 2B, middle), suggesting a trend toward fewer but longer microtubules in the processes. The physical properties of the microtubules, indicated by their straightness, did not change (Fig. 2B, right). To determine how hydrogen peroxide affects microtubule length, we plotted the length distribution in a relative frequency histogram (Fig. 2C). The data show that hydrogen peroxide increases mean microtubule length primarily by reducing the proportion of short microtubules (0.5-2.5 µm length) and increasing the number of long microtubules (>8 µm).

**Fig. 2.**
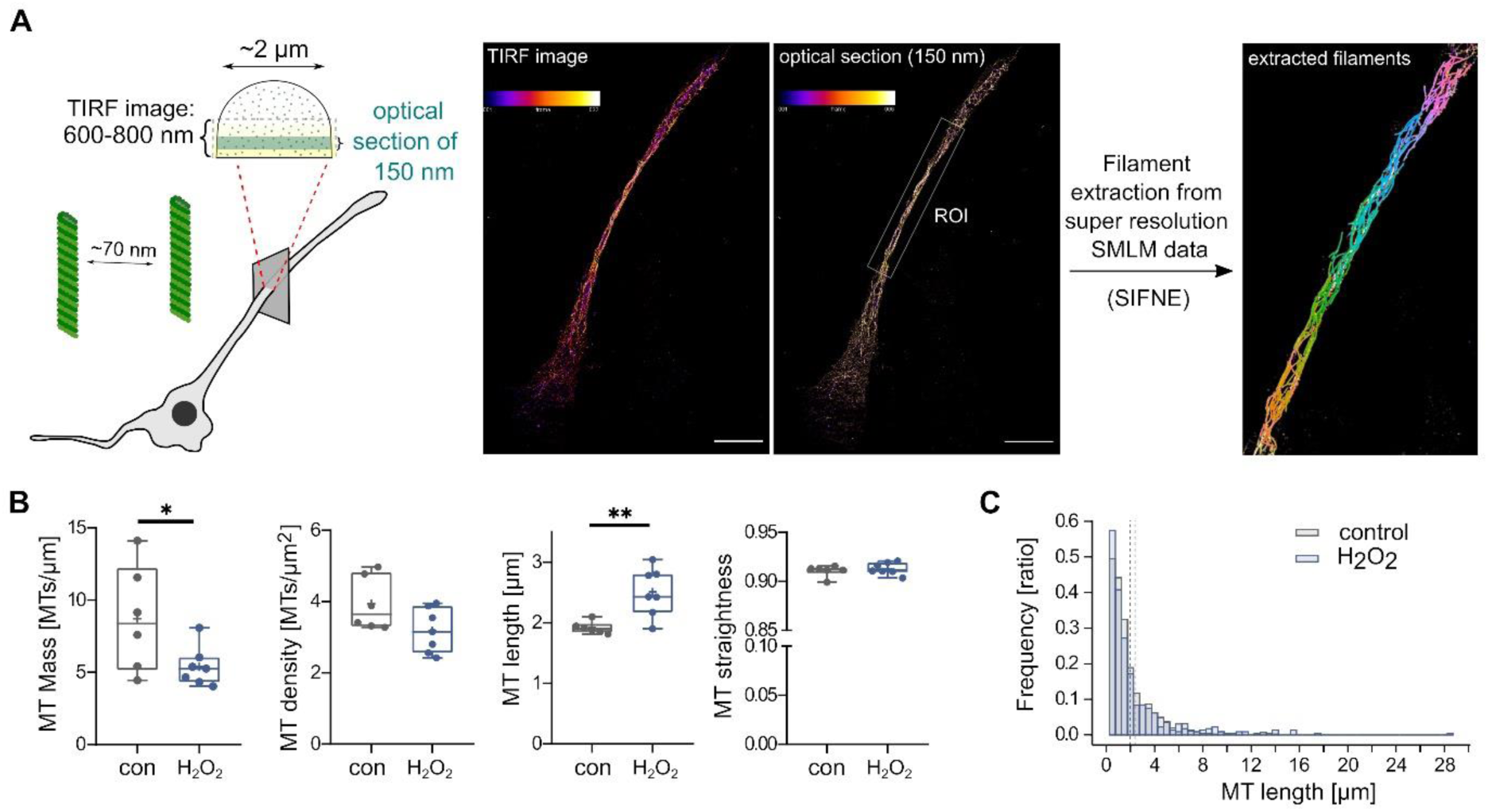
Subtoxic hydrogen peroxide modulates the structure of the axonal microtubule array by reducing microtubule mass and increasing microtubule length. **A.** Shown on the left is a schematic representation of the single molecule localization microscopy (SMLM) approach to quantify MT organization. An indication of the average microtubule spacing in axon-like processes of differentiated PC12 is included. A total internal reflection fluorescence (TIRF) image of the 3D-rendered SML data and a 150 nm optical section with fire color code are shown in the middle. Microtubule filaments as extracted from a selected ROI using SIFNE (single-molecule localization microscopy image filament network extractor) are shown in the rainbow color code on the right. Scale bar 10µm. **B.** Boxplots showing the mass, density, mean length and straightness of microtubules. Each data dot represents the average of a single cell (control, 6 cells with n=902 individual microtubules; H2O2, 7 cells with n=457 individual microtubules). H2O2 treatment reduces microtubule mass but increases mean microtubule length. Statistically significant differences determined by unpaired two-tailed Student’s t-tests are indicated. *p<0.05, **p<0.01. **C.** Distribution of microtubule lengths in a relative abundance histogram. Dotted lines show mean microtubule length under control conditions and with H2O2. The bin width was set to 0.5 µm and the area of the histogram bars is 1.0. H2O2 shifts the length distribution by reducing the proportion of short microtubules (0.5-2.5 µm length) and increasing the number of long microtubules (>8 µm).

Thus, the data indicate that hydrogen peroxide alters the organization of microtubule arrays in axon-like processes by reducing their mass and increasing their length, but does not affect their physical properties.

### Hydrogen peroxide-induced changes in the microtubule array reduce the proportion of mobile vesicles but have no effect on the speed and velocity of axonal transport

A change in axonal microtubule arrangement may have a direct impact on the properties of axonal transport, as C. elegans motor neurons have been shown to pause axonal transport at the ends of microtubules before switching to a new polymer (Yogev et al., 2016). This suggests that altering microtubule tracks can affect the efficiency of axonal transport.

To determine whether the hydrogen peroxide-induced change in the microtubule array affects axonal transport parameters, we used single-vesicle tracking of eGFP-tagged amyloid precursor protein (APP), a key axonal transport cargo (Morotz et al., 2019) (Fig. 3A). Hydrogen peroxide treatment reduced the proportion of mobile vesicles by approximately 20% compared to a control (Fig. 3B, left), indicating a reduction in the total amount of cargo transported. To quantify the movement of mobile vesicles, we determined the effect of hydrogen peroxide on velocity (displacement per time) and speed (run length per time) from the trajectories of each cell analyzed (Fig 3B, middle and right). Both the velocity and speed of APP-vesicles were the same under both conditions, indicating that the change in microtubule length distribution had no effect on the transport of the vesicles on the microtubule tracks once they were moving.

**Fig. 3.**
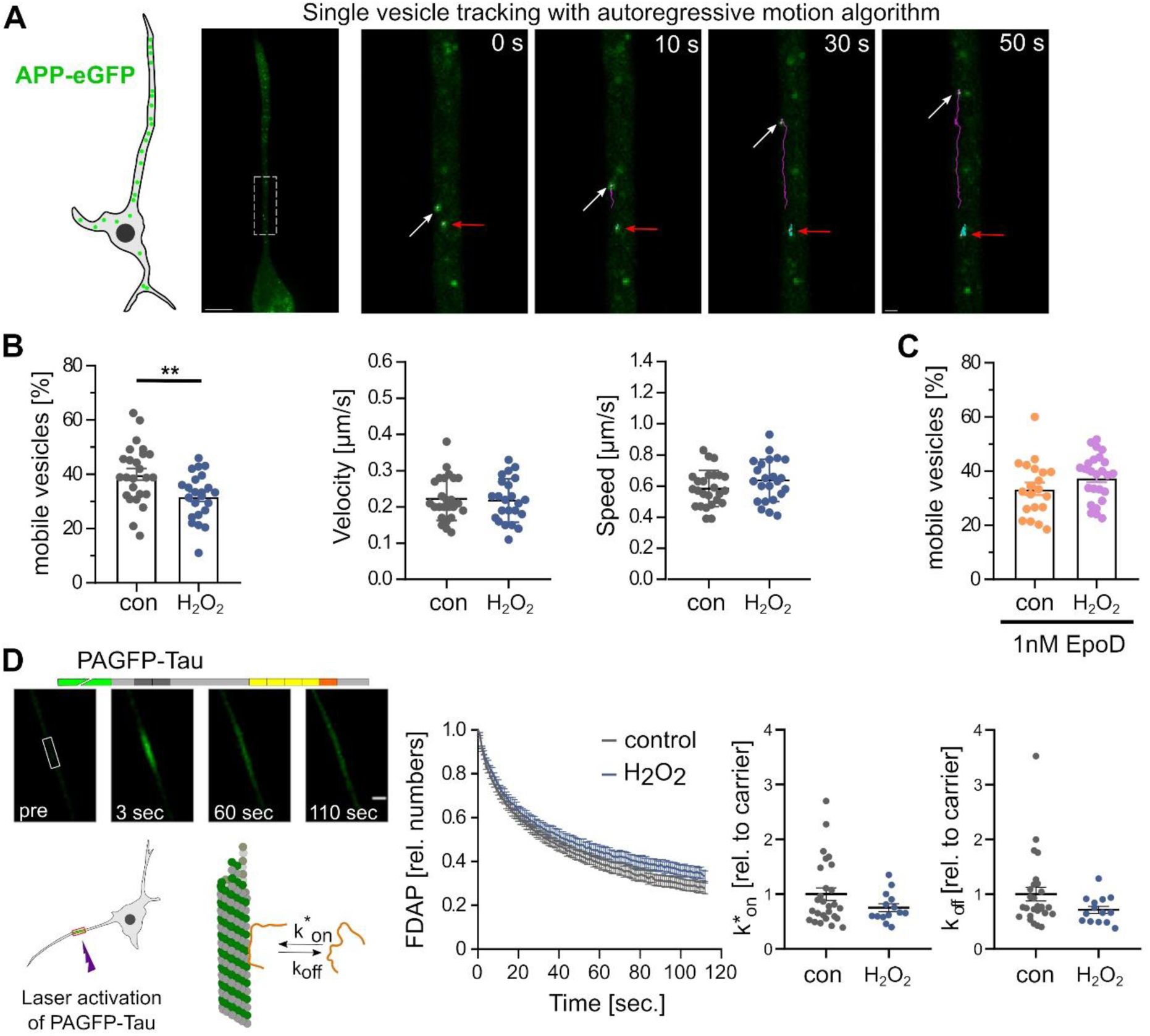
Hydrogen peroxide-induced changes in the microtubule array reduce the proportion of mobile vesicles but have no effect on the speed and velocity of axonal transport, or the interaction with the axonal tau protein. **A.** Tracking APP vesicles in a PC12 cell process using eGFP-tagged APP and an autoregressive motion algorithm. The first micrograph shows an overview of a cell expressing APP-eGFP. The images on the right show selected times of the APP vesicle movement from the part of the process marked by the white box in the overview image. Arrows point to a moving (white) and a stationary vesicle (red). Scale bars, 10 μm (overview) and 1 μm (time lapse). **B.** The bar graph shows proportions of mobile vesicles in the axon-like process under control conditions and in response to H2O2. Velocity and speed of mobile vesicles are shown in the scatter plots on the right (mean ± SEM of n = 25 cells with 1595 trajectories (control) and 23 cells with 1209 trajectories (H2O2); each point represents an average value for one analyzed cell). Statistically significant differences determined by unpaired two-tailed Student’s t-tests are indicated. **p < 0.01. **C.** Proportion of mobile vesicles in cells pretreated with 1 nM EpoD according to the timeline shown in Fig 2D. Each data point represents an average value for a corresponding cell (mean ± SEM of 20 cells with 1276 trajectories (control) and 25 cells with 1720 trajectories (H2O2)). **D.** Effect of H2O2 on the interaction of tau with microtubules. A schematic representation of the FDAP approach and the expressed tau construct is shown on the left. FDAP plots after photoactivation of PAGFP-Tau expressing cells (middle) show a similar fluorescence decay with and without H2O2. Likewise, the scatterplots of the association (k^*^) and dissociation rate constants (koff) (right) show no statistically significant differences. Mean values ± SEM of 25 (control) and 14 (H2O2) cells are shown.

To confirm that the observed reduction in the proportion of mobile vesicles is due to the change in the organization of the microtubule array, we determined the effect of the microtubule stabilizer EpoD. We previously observed that nanomolar concentrations of EpoD shifted the length distribution of microtubules in axon-like processes toward shorter and denser microtubules (Conze et al., 2022b). Indeed, EpoD abolished the effect of hydrogen peroxide on reducing the fraction of mobile vesicles (Fig. 3C), suggesting that the effect was caused by the altered organization of the microtubule array.

Thus, the data indicate that hydrogen peroxide has the potential to influence axonal transport by reducing the total amount of cargo transported, likely due to its effect on modulating the microtubule array.

### Hydrogen peroxide-induced changes in the microtubule array do not affect the dynamic interaction of the axonal tau protein with microtubules

Tau protein is an axonally enriched microtubule-associated protein (MAP) that is thought to regulate axonal microtubule polymerization. Pathological changes in tau are involved in a class of neurodegenerative diseases collectively referred to as tauopathies (Arendt et al., 2016). Under physiological conditions, tau dynamically interacts with microtubules and exhibits a “kiss and hop” behavior that is likely required to regulate microtubule polymerization without affecting the efficiency of axonal transport (Janning et al., 2014). Indeed, post-translational modifications of tau that make tau microtubule interaction less dynamic can lead to axonal transport defects that cause dendritic atrophy in tauopathies (Bakota and Brandt, 2024; Conze et al., 2022a).

Therefore, hydrogen peroxide could induce changes in the interaction of tau with axonal microtubules, which may affect microtubule dynamics, cause the observed reduction in microtubule polymer, and impair microtubule-dependent cargo transport. We used FDAP experiments to determine a possible change in the dynamics of tau interaction with microtubules in axon-like processes of the model neurons. Tau was N-terminally tagged with photoactivatable GFP (PAGFP) and expressed exogenously in PC12 cells, which were differentiated to a neuronal phenotype. Following focal activation of tau in hydrogen peroxide-treated or control cells, tau showed similar dissipation from the activation area (Fig. 3D, left). Application of a refined reaction-diffusion model of the tau-microtubule interaction allowed the determination of the binding constants (k^*^on and koff rates) of the tau microtubule-interaction (Igaev et al., 2014). We did not detect any difference in either kinetic constant as a result of hydrogen peroxide treatment (Fig. 3D, right).

Thus, the data indicate that hydrogen peroxide does not affect the dynamics of tau-microtubule interaction in axon-like processes. Consequently, the data also indicates that the increase in microtubule dynamics as a result of hydrogen peroxide treatment is not caused by a change in the interaction of tau with microtubules.

### Subtoxic hydrogen peroxide modulates the phosphorylation state of components of the microtubule system

Hydrogen peroxide, as a second messenger molecule, is thought to link redox biology to intrinsic signalling pathways (Sies, 2017). Thus, it is likely that subtoxic hydrogen peroxide shapes the axonal microtubule array by affecting the expression or phosphorylation of proteins of the microtubule system. To identify the respective target molecules and their modification, we performed proteomics and phosphoproteomics analysis of differentiated model neurons treated with hydrogen peroxide compared to control conditions (Fig. 4A). GO-term analysis for differentially expressed proteins revealed mainly NADPH-binding and RNA polymerase-binding proteins (Fig. 4B). The only microtubule-related protein with differential expression was MAPRE3 (Microtubule-associated protein RP/EB family member 3, EB3), which showed slightly increased expression in hydrogen peroxide-exposed cells (Fig. 4C). EB3 is known to be a microtubule plus-end binding protein (Akhmanova and Steinmetz, 2015) and promotes microtubule growth by suppressing catastrophe (Komarova et al., 2009). However, it was recently shown to be also relevant to microtubule minus-end organization (Yang et al., 2017). Therefore, the increase in mean microtubule length which we observed in the microtubule array could be due to the increased EB3 expression.

**Fig. 4.**
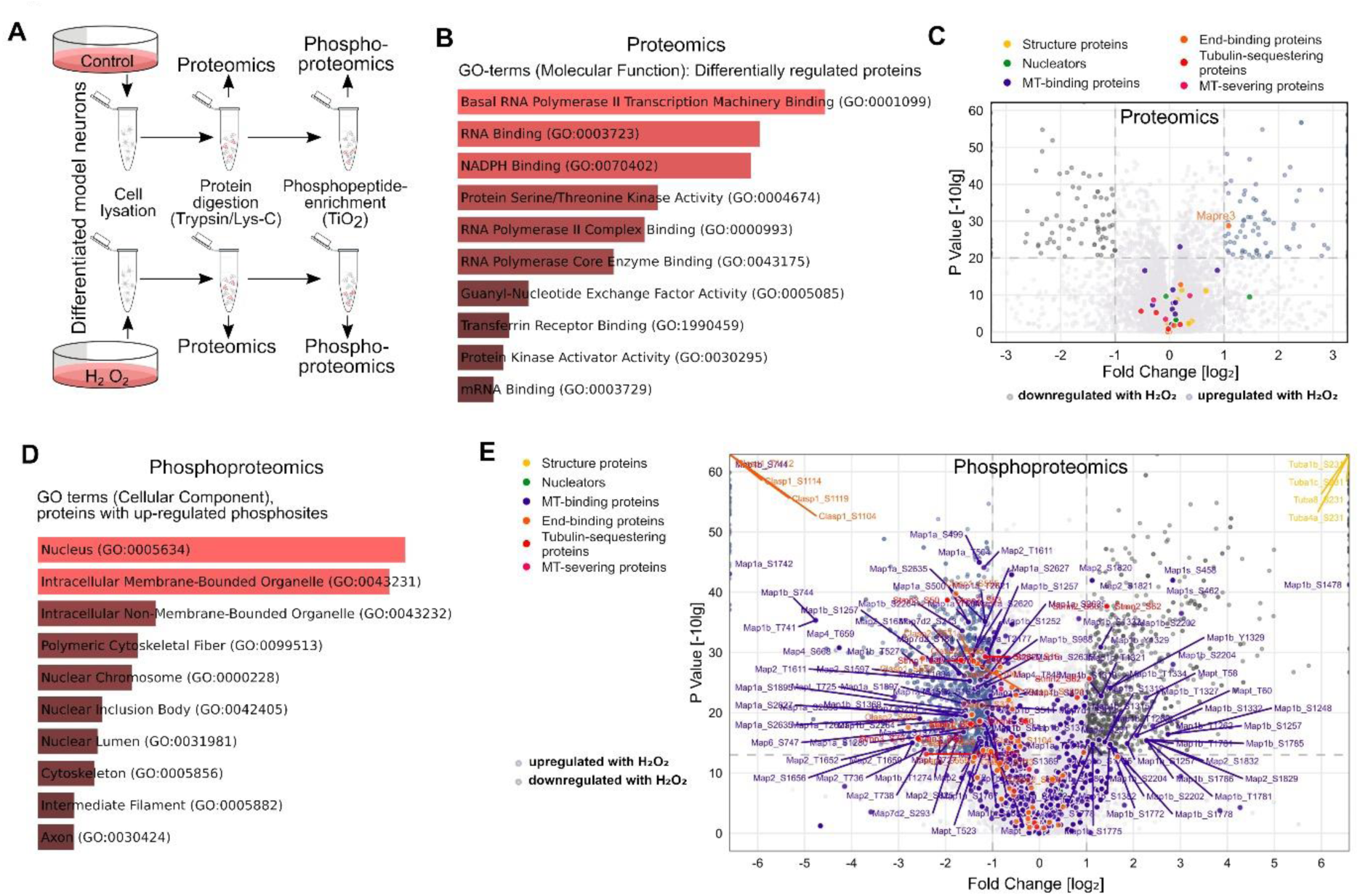
Subtoxic hydrogen peroxide modulates the phosphorylation state of components of the microtubule system. **A.** Schematic representation of the approach for proteomics and phosphoproteomics analyses of differentiated model neurons treated with hydrogen peroxide compared to control. **B.** Bar plot with the top enriched GO-terms for molecular function of all differentially regulated proteins in response to H2O2. The length of the bar indicates the p-value, reflecting the enrichment significance of each term. Note an enrichment of proteins mostly involved in RNA processing and redox modulation. **C.** Volcano plot showing up- and down-regulated proteins in hydrogen peroxide-treated cultures compared to controls. Log2 fold changes are plotted against -log10 p-values. Significant upregulation upon H2O2 treatment is shown in blue, downregulation in dark grey. Members of different groups of microtubule-regulating proteins are indicated. The axes are cut for representation purposes (x-axis from -3 to 3, y-axis from 0 to 60); all the proteins above the limits are shown as points at each limit border. Members of different groups of microtubule-regulating proteins are indicated in different colors, with significantly changed ones labelled with their gene names. Only one protein of the more than 30 identified proteins of the microtubule system (MAPRE3) shows a slight but significant upregulation. **D.** Bar plot with the top enriched GO-terms for cellular components of all proteins with upregulated phosphosites in response to H2O2. The length of the bar indicates the p-value, reflecting the enrichment significance of each term. Note enrichment of nuclear components, organelles and proteins of the cytoskeleton. **E.** Volcano plot showing up- and down-regulated phosphosites in hydrogen peroxide-treated cultures compared to controls. The axes are cut for representation purposes and all phosphosites above the limits are shown as points at each limit border. Coloring and labelling as in C.

In contrast to the moderate effect on the expression of microtubule system proteins, hydrogen peroxide had a strong effect on the phosphorylation of various microtubule proteins, as demonstrated by phosphoproteomic analysis (Fig. 4D, E). GO-term analysis of proteins with upregulated phosphosites revealed the presence of many cytoskeletal proteins, particularly components of the microtubule skeleton (Fig. 4D). Further examination of the proteins with upregulated phosphosites revealed that various functional subgroups of the microtubule system, including structural proteins, tubulin-sequestering proteins and microtubule-associated proteins, exhibited increased phosphorylation (Fig. 4E).

Thus, the data indicate that hydrogen peroxide shapes the axonal microtubule array primarily by modulating the phosphorylation status of different functional groups of the microtubule system.

### Hydrogen peroxide and ROS-generation by arsenite induce differential phosphorylation of MAP1B as the major target of microtubule proteins

If hydrogen peroxide acts as a second messenger molecule, it would be expected to selectively regulate signalling pathways under eustress conditions that differ from the action of other redox modulators, particularly when compared to conditions that induce oxidative distress. Therefore, we decided to compare the effect of subtoxic hydrogen peroxide with ROS-generation by arsenite, a naturally occurring toxicant (Huber et al., 2022).

Phosphoproteomic analysis showed that arsenite also affected the phosphorylation of several functional groups of the microtubule system, including structural proteins, microtubule-binding proteins, microtubule nucleators, and tubulin-binding proteins (Fig. 5A). In fact, arsenite induced a higher number of upregulated phosphosites than hydrogen peroxide, which was more than twice as high for the microtubule system proteins (Fig. 5B). Notably, the overlap of upregulated phosphosites induced by hydrogen peroxide and arsenite was small, and less than 5% (4 out of 82) of the phosphosites increased by arsenite were also increased in the presence of hydrogen peroxide.

**Fig. 5.**
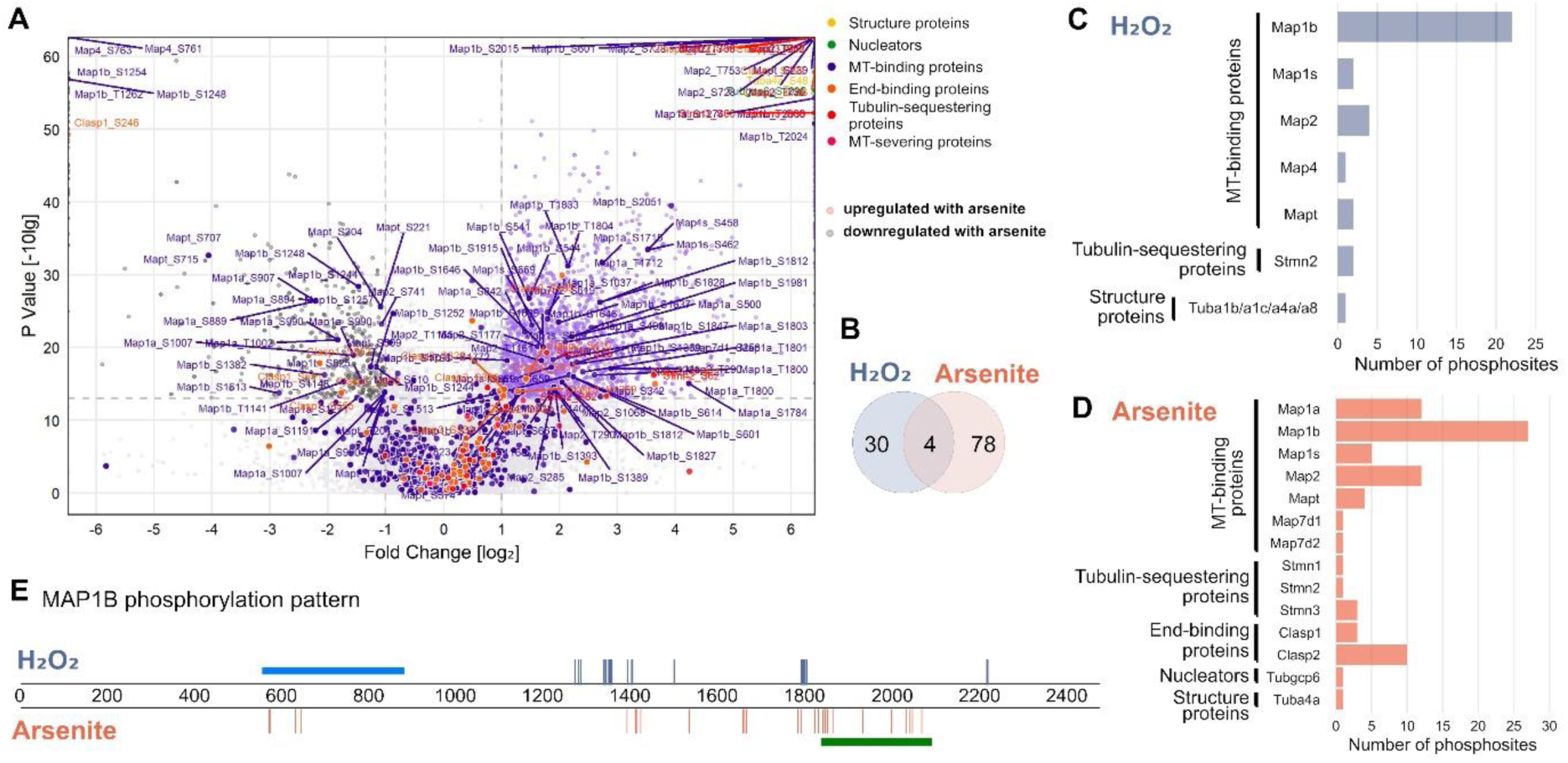
Subtoxic hydrogen peroxide and ROS-generation by arsenite induce differential phosphorylation of MAP1B as the major target of microtubule proteins. **A.** Volcano plot showing upregulated phosphosites in arsenite-treated cells compared to control. Members of different groups of microtubule-regulating proteins are indicated by the same color code as in Fig. 5C. Log2 fold changes are plotted against -log10 p-values. Significant upregulation upon arsenite treatment is shown in orange, downregulation in dark grey. The axes are cut for representation purposes and all phosphosites above the limits are shown as points at each limit border. **B.** Venn diagram showing low overlap of phosphosites of microtubule- regulating proteins upregulated in response to hydrogen peroxide or arsenite. **C, D.** Distribution of upregulated phosphosites on different proteins of the microtubule system in response to hydrogen peroxide (C) or arsenite (D). Note that MT-binding proteins and especially MAP1B are the main target. **E.** Graphical representation of the different phosphosites on MAP1B that are upregulated in response to hydrogen peroxide or arsenite. The blue bar shows the microtubule-binding region according to the deletion study by (Noble et al., 1989). Phosphorylated epitopes recognized by the monoclonal antibody SMI-31 (Johnstone et al., 1997), which detects disease-associated mode I phosphorylation sites, which cause a loss of the microtubule stabilizing activity, are indicated by the dark green bar.

A comparison of the target proteins that showed increased phosphorylation revealed that twice as many proteins of the microtubule system were altered in arsenite-exposed cells (Fig. 5C, D). In both cases, the microtubule-binding protein MAP1B was the main target. MAP1B is predominantly expressed in the nervous system and has been implicated in the regulation of axonal elongation and guidance (Bouquet et al., 2004; Gonzalez-Billault et al., 2001). MAP1B showed a complex pattern of increased phosphorylation sites with both hydrogen peroxide (22 sites) and arsenite (27 sites) (Fig. 5E). Notably, only one of the sites (S1772) showed increased phosphorylation under both conditions. This specificity of increased phosphorylation at selected sites was also present in other MAPs such as tau (total of 6 phosphosites, no overlap) and MAP2 (total of 16 phosphosites, no overlap).

Thus, the data indicate that hydrogen peroxide leads to a specific phosphorylation pattern of proteins of the microtubule system that is largely distinct from the effects of arsenite as a toxic redox modulator, reflecting conditions of oxidative distress. The data also point to MAP1B as the main target for both redox modulators, suggesting that MAP1B is the master regulator of redox-mediated axonal microtubule dynamics.

### Cell-wide phosphoproteome analysis of predicted upstream kinases in neuronally diffentiated cells reveals a pattern of inversely regulated kinases by hydrogen peroxide and arsenite

The fact that the two redox modulators hydrogen peroxide and arsenite each have a different effect on the phosphorylation of microtubule-regulating proteins suggests that oxidative eustress and distress influence specific signalling cascades differently. Therefore, we performed a kinase enrichment analysis using all changed phosphosites as input to predict the upstream kinases responsible for the different phosphorylation events observed (Kuleshov et al., 2021). Remarkably, we observed no overlap between the predicted upstream kinases that were activated by the two redox modulators (Fig. 6A). Prominent upstream kinases activated by hydrogen peroxide included kinases known to suppress apoptosis such as DNA-dependent protein kinase PRKDC (Yue et al., 2020) or the casein kinase 2 subunit CSNK2A2 (Ahmad et al., 2008). Other kinases are known to be involved in regulating brain metabolism such as the pyruvate dehydrogenase kinases PDK2 and 3 (Wang et al., 2023). On the other hand, prominent predicted upstream kinases activated by arsenite were several members of the MAP kinase signal transduction pathway (MAPK3, MAP3K4, MAP2K1, 3, 4, 6, and 7), mammalian target of rapamycin (mTOR), and serum/glucocorticoid regulated protein kinases SGK1, 2 and 3. Deregulation of the MAPK signaling and mTOR pathway has been implicated in the development of several neurodegenerative diseases (Ahmed et al., 2020; Perluigi et al., 2015) and upregulation of SKG is associated with various neurodegenerative disorders (Kwon et al., 2021).

**Fig. 6.**
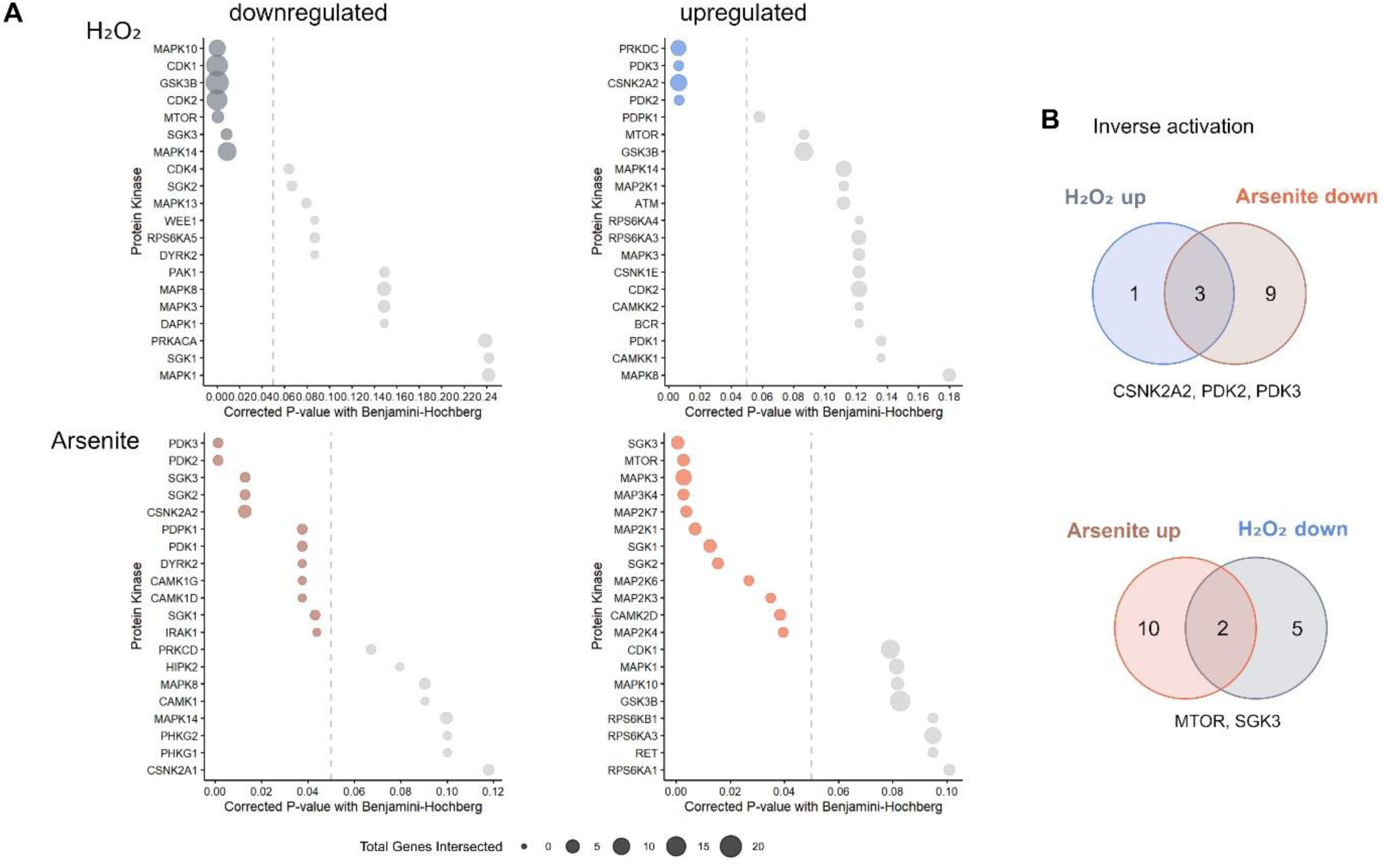
Cell-wide phosphoproteome analysis of predicted upstream kinases in neuronally differentiated cells reveals a pattern of inversely regulated kinases by hydrogen peroxide and arsenite. **A.** Kinase enrichment analysis of the phosphoproteomics data to identify the pattern of kinases responsible for increased phosphorylation of all cellular proteins in response to hydrogen peroxide (left) and arsenite (right). Note that there is no overlap between significantly upregulated upstream kinases with hydrogen peroxide and arsenite. **B.** Venn diagrams showing upstream kinases leading to a reverse change in phosphorylation, i.e. induction of increased phosphorylation with hydrogen peroxide and reduced phosphorylation with arsenite (top) and increased phosphorylation with arsenite and reduced phosphorylation with hydrogen peroxide (bottom). The respective upstream kinases with reverse change are indicated below the corresponding Venn diagram.

Of particular importance for the regulation of differential phosphorylation by certain redox modulators could be those signalling pathways that lead to a reverse change in phosphorylation, i.e. to activation under one condition, but to inhibition under the other. The respective activated upstream kinases for hydrogen peroxide versus arsenite include pyruvate dehydrogenase kinases (PDKs), which play a crucial role in aerobic metabolism, linking glycolysis to the tricarboxylic acid cycle and ATP generation (Wang et al., 2021), and a subunit of casein kinase 1, a major contributor to the generation of the human phospho-proteome (Borgo et al., 2021) (Fig. 6B). In turn, the upstream kinases for arsenite versus hydrogen peroxide include mTOR and glucocorticoid-regulated kinase 3 (SGK3), kinases whose dysregulation has been linked to metabolic dysfunction and disease (Liao et al., 2022; Maiese, 2020).

Taken together, cell-wide phosphoproteome analysis shows that different signalling pathways are mutually exclusively activated by oxidative eustress and distress. While treatment with subtoxic hydrogen peroxide (reflecting oxidative eustress) is mainly associated with kinases suppressing apoptosis and regulating brain metabolism (PRKDC and CK2) and cellular metabolism (PDKs), treatment with the toxic redox modulator arsenite (reflecting oxidative distress) resulted in activation of signalling pathways associated with neurodegeneration (mTOR, SGKs). The different phosphorylation patterns of microtubule-regulating proteins, in particular MAP1B, is reflected by a mutually exclusive activation of signaling pathways at eustress versus distress conditions.

## DISCUSSION

Reactive oxygen species are involved in a variety of physiological cellular functions such as cell proliferation, differentiation and maturation (Beckhauser et al., 2016). However, when ROS accumulation exceeds antioxidant defense mechanisms, it leads to oxidative distress and promotes pathological conditions in the brain (Pratico et al., 2002). The development and maturation of neurons critically depends on the microtubule cytoskeleton. Pathological changes in the microtubule system and microtubule-dependent functions such as axonal transport occur early in the disease process and disruption of microtubule dynamics may be a key mechanism contributing to neurodegeneration (Penazzi et al., 2016a). However, it is largely unknown how physiological, nontoxic ROS levels (oxidative eustress) influence axonal microtubule dynamics and microtubule-dependent functions and how they differ from conditions that trigger oxidative distress.

Here, we show that (1) a subtoxic concentration of the redox messenger hydrogen peroxide increases microtubule dynamics and shapes microtubule organization in axon-like processes, that (2) hydrogen peroxide modulates the phosphorylation state of different functional groups of the microtubule system, and that (3) subtoxic hydrogen peroxide induces a complex and ROS-specific phosphorylation pattern of MAP1B as the main target protein of the microtubular system. By cell-wide phosphoproteome analysis, we further provide evidence that (4) different signalling pathways are mutually exclusive activated by oxidative eustress and distress, reflecting the distinct phosphorylation patterns of microtubule-related proteins. While oxidative eustress is associated with the activation of pathways related to kinases that suppress apoptosis and regulate brain metabolism (PRKDC, CK2, PDKs), treatment with the toxic redox modulator arsenite led to the activation of signalling pathways related to neurodegeneration (mTOR, SGKs).

Axonal microtubules exhibit a unique organization in that they exist in relatively short fragments with a uniform orientation and their dynamic plus-ends pointing towards the axon tip. Although most axonal microtubules are more stable than the microtubules of dendrites, axons also possess dynamic microtubules that undergo phases of polymerization and depolymerization, a process known as dynamic instability (Baas et al., 2016). The dynamic microtubules in the axon can act as sensors of the cellular microenvironment and enable the rapid reorganization of the cytoskeleton in response to changes in the environment (Coles and Bradke, 2015). The regulation of microtubule nucleation and the dynamics of their polymerization could therefore play an important role in local axon homeostasis for axon maintenance, function and pathology (Hahn et al., 2019). Regulation by ROS can thus couple neuronal activity to the local organization of the microtubule array. High neuronal activity would lead to an increase in ROS species due to more active mitochondria. A physiologically increased hydrogen peroxide content would then lead to a rearrangement of the microtubule array by decreasing microtubule density and increasing mean microtubule length, accompanied by a reduced amount of transported cargo, as we observed in our experiments (Fig. 7). This would create a negative feedback loop to dampen over-activation of the neuron.

**Fig. 7.**
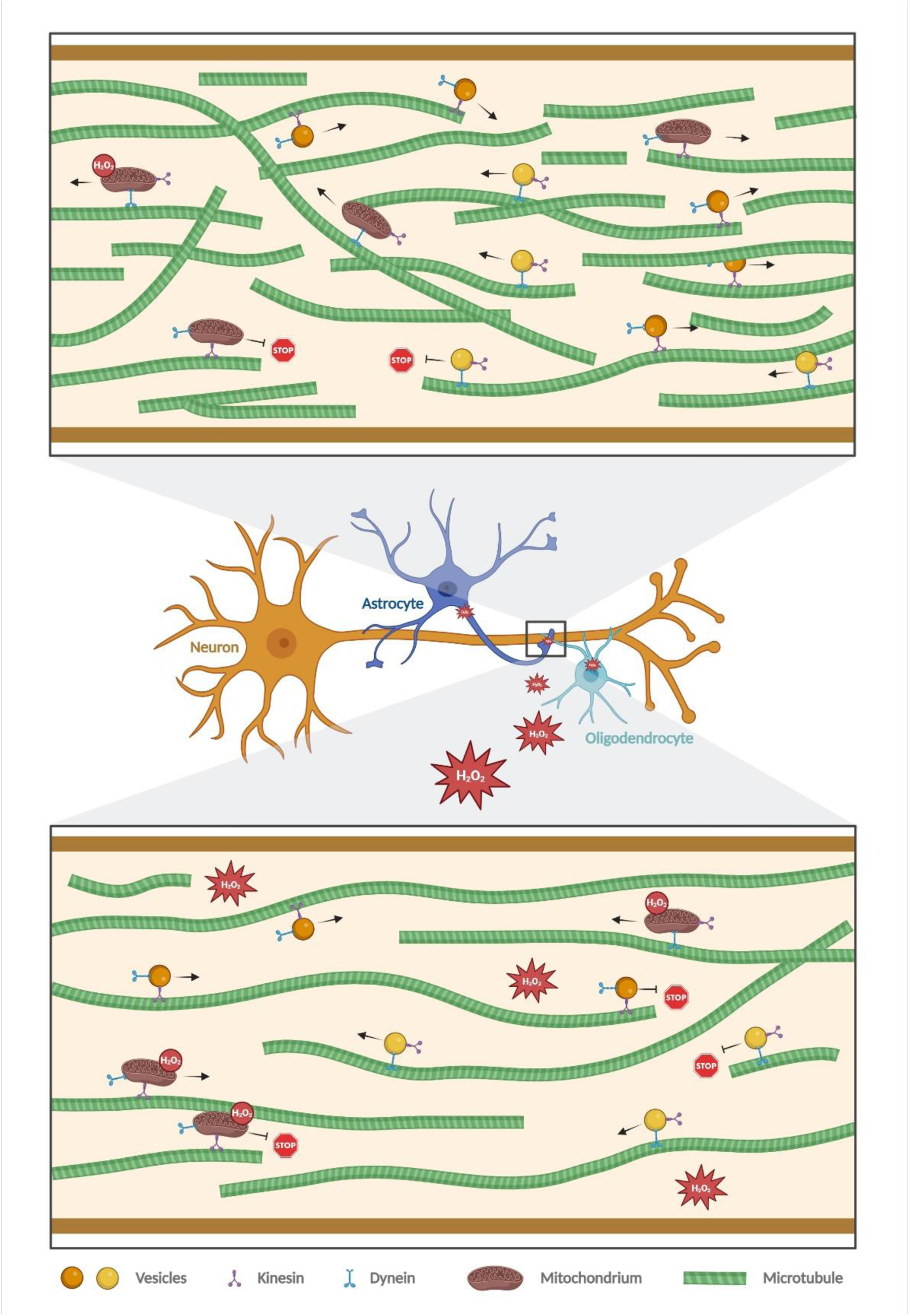
Schematic representation of the effect of hydrogen peroxide on axonal microtubule organization and microtubule-dependent transport. Hydrogen peroxide diffuses through cells and tissues and can reach neighboring axons when produced in oligodendrocytes or astrocytes. In addition, it is also produced by mitochondria in the axons. In axons, H2O2 causes a reorganization of the microtubule cytoskeleton toward longer but less dense microtubules, which reduces the efficiency of vesicle transport because it becomes more difficult for vesicles to change the microtubule track. Figure created with BioRender.com.

Hydrogen peroxide may also play a role in communication between glial cells and neurons. As a messenger molecule, hydrogen peroxide diffuses through cells and tissues, which would allow myelinating oligodendrocytes, which are known to provide metabolic and functional support to the underlying axon (Duncan et al., 2021; Simons and Nave, 2015), to modulate the structure and dynamics of axonal microtubules. Oligodendrocytes, in turn, could also act as a sink for hydrogen peroxide produced locally in the axon. Therefore, it would be interesting to determine the possible crosstalk of oligodendrocytes and neurons with respect to redox signalling.

Our data show that several functional groups of the microtubule system are differentially phosphorylated due to hydrogen peroxide signalling. By far the largest group is microtubule-associated proteins (MAPs), followed by the tubulin sequestering protein stathmin 2. This suggests that the other groups of microtubule-regulating proteins such as microtubule-severing factors, end-binding proteins or microtubule nucleators are not involved in the immediate response to hydrogen peroxide with regard to axonal microtubule homeostasis. Interestingly, microtubule-binding proteins and tubulin-sequestering proteins were also identified as the most important drivers for the development of increased neuronal complexity during vertebrate evolution, as they showed the largest increase in the number of orthologs and predicted protein-coding splice variants (Trushina et al., 2019b). Thus, MAPs and tubulin-sequestering factors appear to be the most important microtubule-regulating proteins in adapting the microtubule skeleton to changes in the environment, both on an evolutionary scale and in terms of regulating axonal microtubule homeostasis. Under oxidative distress conditions (arsenite treatment), end-binding proteins and a microtubule nucleator also joined the functional groups modified by phosphorylation, which may indicate that increased phosphorylation of end-binding proteins in particular could be responsible for the pathological changes of the axonal microtubule array.

Our data show that MAP1B is the main target of phosphorylation in both eustress and distress conditions and approximately two third of the phosphorylation sites of microtubule-related proteins that are upregulated in eustress conditions belong to MAP1B. MAP1B appears to preferentially associate with tyrosinated (dynamic) microtubules, rather than detyrosinated (stable) microtubules (Tymanskyj et al., 2012), which may increase the pool of dynamic microtubules (Tortosa et al., 2013; Utreras et al., 2008). Such behavior could be important for the local regulation of axonal microtubule dynamics and polymerization. Phosphorylation of MAP1B at different sites can regulate the local fine-tuning of neurite branching and microtubule dynamics (Barnat et al., 2016; Scales et al., 2009; Ulloa et al., 1993). Historically, two types of MAP1B phosphorylation have been distinguished. Mode I phosphorylation induces a shift in electrophoretic mobility and decreases with development, while mode II phosphorylation does not affect electrophoretic mobility and remains unchanged (Kawauchi et al., 2005). Phosphorylation at mode I sites results in a loss of microtubule-stabilizing ability (Goold et al., 1999) and mode I phosphorylated MAP1B is also observed in neurofibrillary tangles (NFTs) and dystrophic neurites in Alzheimer’s disease brains (Ulloa et al., 1994). This suggests that mode I phosphorylation may be associated with nervous system pathogenesis and reflects the MAP1B phosphorylation at oxidative distress conditions. In contrast, mode II phosphorylation is mediated for example by casein kinase 2 (Ulloa et al., 1993), a predicted upstream kinase with inverse regulation with hydrogen peroxide and arsenite. Regions of phosphorylated epitopes recognized by the monoclonal antibody SMI-31 implicated in detecting mode I phosphorylation sites (Johnstone et al., 1997) are present both after arsenite treatment (1836-2076) and hydrogen peroxide treatment (1244-1264) (Fig. 6E). It will be interesting to determine the functional consequences of the hydrogen peroxide-induced change in phosphorylation compared to the arsenite-induced change, in order to determine sites that may be crucial for the physiological versus pathological regulation of microtubule polymerization. However, the remarkable complexity of MAP1B phosphorylation makes this a difficult undertaking.

Increased phosphorylation of the microtubule-associated protein tau is associated with the development of tauopathies such as Alzheimer’s disease (Arendt et al., 2016; Trushina et al., 2019a). Remarkably, our data show that tau is not a major target in either oxidative eustress or distress. This is consistent with our observation that treatment with hydrogen peroxide does not affect the tau-microtubule interaction, as many of the disease-related tau phosphorylation events negatively affect tau binding to microtubules. Thus, the data indicate that increased phosphorylation of tau is not involved in the regulation of microtubule polymerization under oxidative eustress conditions and that increased phosphorylation of tau and the concomitant dysregulation of axonal microtubule polymerization is a later event during neurodegeneration.

## ACKNOWLEDGMENTS

We thank Maxim Skulachev (Mitotech S.A., Luxembourg) for providing SkQ1 and TPP, and Amos Smith 3^rd^ (University of Pennsylvania) for providing EpoD. This work was supported by the Deutsche Forschungsgemeinschaft (DFG BR1192/14-1 to RB).

## AUTHOR CONTRIBUTIONS

Christian Conze: Investigation, Writing - Original Draft; Nataliya I. Trushina: Investigation, Writing - Original Draft; Nanci Monteiro-Abreu: Investigation, Writing - Original Draft; Daniel Villar Romero: Investigation, Writing - Original Draft; Eike Wienbeuker: Investigation, Writing - Original Draft; Anna-Sophie Schwarze: Investigation; Michael Holtmannspötter: Investigation, Writing - Original Draft; Lidia Bakota: Writing - Original Draft, Supervision; Roland Brandt: Conceptualization, Writing - Original Draft, Supervision, Funding acquisition.

The authors declare that no generative AI or AI-assisted technologies were used in the preparation of this manuscript.

